# Chaphamaparvovirus Antigen and Nucleic Acids are not Detected in Kidney Tissues from Cats with Chronic Renal Disease or Immunosuppressive Diseases

**DOI:** 10.1101/2021.03.30.437777

**Authors:** AO Michel, TA Donovan, B Roediger, Q Lee, C Jolly, S Monette

**Author notes:** Corresponding author: Sebastien Monette, Laboratory of Comparative Pathology, Center of Comparative Medicine and Pathology, Memorial Sloan Kettering Cancer Center, The Rockefeller University, Weill Cornell Medicine, New York, NY 10065, USA.

## Abstract

Mouse Kidney Parvovirus (MKPV) was recently recognized as the cause of murine inclusion body nephropathy, a disease reported for over 40 years in laboratory mice. Immunodeficient mice are persistently infected with MKPV, leading to chronic renal disease, morbidity and mortality whereas immunocompetent mice seroconvert with mild renal pathology. Given the high incidence of MKPV infection in wild mice in the New York City area, the first goal of this study was to evaluate the possibility of MKPV involvement in feline chronic kidney disease (CKD) from the same geographic region. As MKPV and related parvoviruses recently described in other animal species appear to have a tropism for kidney tissue, the second goal was to investigate the possible role of a virus of this group, other than MKPV, in the development of feline CKD, Presence of MKPV and related viruses was investigated in feline renal samples using PCR, RNA *in situ* hybridization (ISH) and immunohistochemistry (IHC). Cats were divided into three groups: normal (N=25), CKD (N=25) and immune suppressed (N=25). None of the kidney tissues from any of the 75 cats revealed the presence of MKPV DNA, RNA or antigen expression. Nor was “fechavirus” detected using PCR in renal tissue from cats with chronic kidney disease. We conclude that MKPV is an unlikely cause or contributor to feline CKD.

## Introduction

After eluding researchers for decades, the etiology of inclusion body nephropathy (IBN) in laboratory mice was recently determined to be a novel virus named mouse kidney parvovirus (MKPV).^1,10,14,15^ Naturally occurring morbidity and mortality due to chronic renal disease attributed to IBN has been documented in laboratory mouse strains lacking functional B and T cells (*Rag1*-knockout, scid and NOD scid gamma (NSG) strains) at multiple institutions.^6,14^ IBN in these mice is a chronic progressive disease characterized by tubular degeneration and necrosis with intranuclear inclusion in tubular cells, interstitial fibrosis, azotemia and anemia, progressing over the course of several months and leading to clinical signs and death attributed to renal failure. Milder, subclinical renal lesions associated with similar intranuclear lesions in tubular cells have been described in immunocompetent mice (ICR and SW:Tac stocks), and mice lacking only T cells (athymic nude stocks).^1,14^ After multiple attempts at detecting known viruses in affected kidneys by immunohistochemistry (IHC), polymerase chain reaction (PCR) and electron microscope, metagenomic analysis was performed on nucleic acids extracted from diseased murine kidneys in order to investigate the presence of a putative novel pathogen.^1,10,14,15^ This analysis yielded sequences with homology to nonstructural (NS1) and structural (VP1) parvoviral proteins.^14^ Sequencing and phylogenetic analysis of the conserved regions of the NS1 protein confirmed a novel parvovirus termed MKPV, which was highly divergent from known murine parvoviruses, mouse parvovirus and minute virus of mice, and was most closely related to a virus recently described in common vampire bats in Brazil. Concurrently to, and independently of, the discovery of MKPV in laboratory mice, a virus named murine chapparvovirus (MuCPV) was discovered by metagenomic analysis in wild mice trapped in residential buildings in New York City.^19^ The MKPV sequences from laboratory mice are 98% identical to those or MuCPV from wild mice, and both viruses were subsequently assigned to a single species, *Rodent chaphamaparvovirus* 1 as the type species for the new genus *Chaphamaparvovirus*^12^ PCR survey of anal swabs of wild mice from multiple residential buildings in NYC revealed a prevalence of 13% to 45% for MuCPV infection.^6^

All chaphamaparvoviruses described to date have been detected by metagenomic sequencing, and so far only MKPV and TiPV (from *Tilapia* fish) have clear causal links with clinical signs or lesions in their hosts.^3,6,9,14^ TiPV was propagated *in vitro* then inoculated to induce disease in fulfillment of Koch’s postulate,^9^ while the causal relationship between MKPV and IBN was demonstrated by a series of experiments that fulfilled Fredrichs and Rellman’s criteria.^14^ Experimental horizontal transmission by soiled bedding transfer and co-housing was demonstrated in mice, and the virus was confirmed to be shed in the urine and possibly the feces. Immunohistochemistry using antisera from immunocompetent, C57BL/6 and Tac:SW mice co-housed with MKPV-positive but not naïve control mice showed immunoreactivity with the inclusion bodies of disease affected mice.^14^ Colocalization of MKPV nucleic acids with tubules displaying inclusions and pathologic changes was demonstrated by *in situ* hybridization.^14^ Diagnostic screening of laboratory mouse samples demonstrated that the virus is widely present in research institutions on 3 continents.^6,14^ MKPV is exquisitely kidney-tropic, a closely-related vampire bat *Chaphamaparvovirus* was detectable in kidney but not spleen, and distantly-related TiPV seems to infect kidneys more efficiently than most other *Tilapia* fish tissues.^3,6,9^

Chronic renal disease is a common condition of domestic cats, affecting up to 49% of cats older than 15 years, and is a significant cause of morbidity and mortality.^2^ Many causes have been proposed to contribute to the disease, including diet, genetics, and infectious agents, but to date, no single unifying etiology has been confirmed. Recent studies have associated renal disease with feline morbillivirus or feline immunodeficiency virus, but although they were suggested as etiologic factors in the renal pathology of cats, no direct causal relationship has been established, while other studies associating feline foamy virus and renal disease show conflicting results.^5,13,16,18,20,8^ More recently, a novel chaphamaparvovirus was identified in feces from cats affected by an unexplained outbreak of diarrhea and vomiting in which common enteric viral and bacterial pathogens were not identified, and was named fechavirus, most closely related to a canine chaphamaparvovirus, although no direct disease causality was proven.^7^ At this time, the prevalence of chaphamaparvoviruses in cats and their role in disease is unclear. In mice, infection with MKPV infection results in persistent viral replication in renal tubular epithelial cells, leading to a spectrum of renal lesions and clinical disease that depends on the host immune status. This disease of mice resembles the spectrum of CKD observed in cats clinically and pathologically, with the exception of intranuclear inclusions which are not described in the feline disease. However, intranuclear inclusions are also not a prominent feature of MKPV infection in immune-competent mice.^3^ Given the high PCR prevalence of MKPV/MuCPV in wild mice living in residential buildings and the strong predator-prey interaction between cats and mice, it is probable that cats are frequently exposed to rodent chaphamaparvoviruses via predation. We hypothesized that *Rodent chaphamaparvopvirus* 1 may infect cats and contribute to feline CKD. Alternatively, we hypothesized that fechavirus or an currently unknown chaphamaparvovirus having cats as its natural host may contribute to feline CKD in a manner similar to MKPV in mice. To test both hypotheses, we performed a retrospective study that aimed to assess diseased and non-diseased renal tissues from cats with normal and deficient immune statuses, using methods that detect proteins and nucleic acids of MKPV/MuCPV and related chaphamaparvoviruses.

## Materials and methods

### Cohort selection and histologic grading

Formalin-fixed, paraffin-embedded (FFPE) kidney tissues from 75 cats were obtained from the archives of the Department of Pathology at The Animal Medical Center. Cats were divided into 3 groups (CKD, immune suppressed, control), with 25 cats in each group. Case selection was performed by searching the necropsy database for reports between 2008 and 2018 that contained certain keywords, and for which the clinical history and/or medical records met the inclusion criteria detailed below.

Group 1 (CKD; N=24) inclusion criteria included cats with a histologic diagnosis of chronic renal disease (keywords: chronic tubulointerstitial nephritis and interstitial fibrosis) and azotemia on plasma biochemical profile at least 3 months prior to postmortem examination. Azotemia was defined as blood urea nitrogen (BUN) and/or creatinine (creat) levels above the reference ranges (BUN 16-37 mg/dL, creat 0.9-2.5 mg/dL). Kidneys were histologically graded using a simplified scheme modified from previously published scales.^2,11^ Group 1 cats required cortical inflammation, cortical fibrosis and tubular degeneration with a grade of at least 3 (Table 1). This group contained kidneys graded as mild (8 cats), moderate (8 cats) and severe (8 cats). Excluded cats were those with the following diagnoses: papillary necrosis, renal dysplasia, glomerulonephritis, polycystic kidney disease, obstructive nephropathy, stents or subcutaneous ureteral bypass, nephroliths, bacterial nephritis, amyloidosis and renal neoplasia.

**Table 1:** Grading Scheme for Severity of CKD

| Grade | Cortical Inflammation | Percent Cortical Fibrosis | Cortical Tubular Degeneration | Totals |
| --- | --- | --- | --- | --- |
| 0 | None/Rare | None/Rare | None/Rare |  |
| 1 | Mild, scattered (<25%) | Mild, scattered, multifocal (<25%) | Focal, scattered | 3-4 Mild |
| 2 | Moderate (25-50%) | Moderate (25-50%) | Multifocal to coalescing | 5-6 Moderate |
| 3 | Diffuse or coalescing (>50%) | Diffuse or coalescing (>50%) | Entire nephron units affected | 7-9 Severe |

Grading Scheme Modified from McLeland et al.<sup>10</sup> and Chakrabarti et al.<sup>1</sup>

Group 2 (immune suppressed; N=26) required a clinical or histologic diagnosis of diseases associated primarily with immune suppression, with or without renal disease. The database was searched for the following keywords: Feline infectious peritonitis virus (FIPV), Feline immunodeficiency virus (FIV), Feline Leukemia virus (FeLV), Feline Panleukopenia virus, Feline Herpesvirus (FHV-1) and Cryptococcus. The severity of the renal lesions from cats in group 2 was also graded according to the modified scheme (Table 1). This group included an additional cat that was initially assigned to group 1, but upon further evaluation was discovered to be immunosuppressed, given a diagnosis of cryptococcosis.

Group 3 (control; N=25) cats required bloodwork without azotemia within 3 months of euthanasia, lack of a post-mortem gross diagnosis involving the kidneys and a CKD histologic diagnosis of 0 for inflammation, fibrosis and tubular degeneration in the grading scale (Table 1).

### DNA extraction, PCR reaction and *in silico* PCR reaction

Five 10 μm-thick paraffin scrolls were obtained from each FFPE block and analyzed by PCR to detect the presence of chaphamaparvovirus DNA. Redundant consensus primers (named 961–963) were designed to amplify a 78 bp VP fragment found to be well-conserved in the sequences of MKPV/MuCPV (*Rodent chaphamaparvovirus 1*, GenBank accession MH670588), rat parvovirus 2 (*Rodent chaphamaparvovirus 2*, GenBank accession KX272741) and chaphamaparvoviruses identified from kidney tissue of two non-rodents: *Chiropteran chaphamaparvovirus 1*, identified in *Desmodus rotundus* (vampire bats) in Brazil (GenBank accession KX907333.1) and “CKPV” identified in a *Cebus imitator* (capuchin) in Costa Rica, currently unclassified (GenBank accession MN265364).^6,17^ Because this sequence is imperfectly conserved in the chaphamaparvovirus recently discovered in cats named fechavirus, we then designed a pair of redundant primers (named 974–975) optimized to detect fechavirus DNA (Figure S1; see the online version of the manuscript for supplemental material).

PCR was performed on cats from group 1 (N=24) and one cat from group 2. Methods for DNA extraction are identical as previously reported.^14^

For PCR, 50–100ng of DNA was added to a cocktail comprising Phire Hot Start II DNA Polymerase (Thermo Scientific, Vilnius, Lithuania), 1x Phire Reaction Buffer, 0.2mM dNTP, 0.5μM of forward primer 961 (5’-CARCARYTNGCWATYCAAGG), 0.25μM of reverse primer 962 (5’-ATCRTCTT<u>GTG</u>CTCCTARTG), 0.25μM of reverse primer 963 (5’-ATCRTCTT<u>TGA</u>CTCCTARTG) in a total 20μL reaction volume; note that primers 962 and 963 were identical but for the 3 bases underlined. The cycling conditions were: initial denaturation at 98°C for 30s, followed by 10 cycles of “touch-down” PCR with denaturation at 98°C for 5s, annealing from 60°C to 50°C for 5s (−1°C per cycle), extension at 72°C for 15s, followed by 30 cycles of denaturation at 98°C for 5s, annealing at 50°C for 5s and extension at 72°C for 15s, and concluded with a final extension at 72°C for 5min. PCR reactions to specifically detect fechavirus were identical, but replaced primers (61-963 with 0.5μM of forward primer 974 (5’-CARCAACTAGCAATACAAGG) and 0.5μM of reverse primer 975 (5’-ATCATCTTGTGC TCCTAATG). 4μL of completed PCR products were analyzed using a Fragment Analyzer 5200 fitted with 33 or 55cm electrophoresis capillaries loaded with matrix capable of resolving dsDNA between 35 and 1500bp (Advanced Analytics, Agilent, Santa Clara, CA, USA). Capillary absorbance traces were analyzed and converted into pseudo-gel images using PROSize 3.0 software (Agilent, Santa Clara, CA, USA). To determine PCR sensitivity, two oligonucleotides 971 and 972 corresponding to nucleotides 2836-2922 of accession MN794869 and two nucleotides 2904-2990 of accession MN396757 (Fig 1A), respectively (synthesized by Integrated DNA Technologies, Singapore), were serially diluted to provide calibration templates.

**Figure 1.**
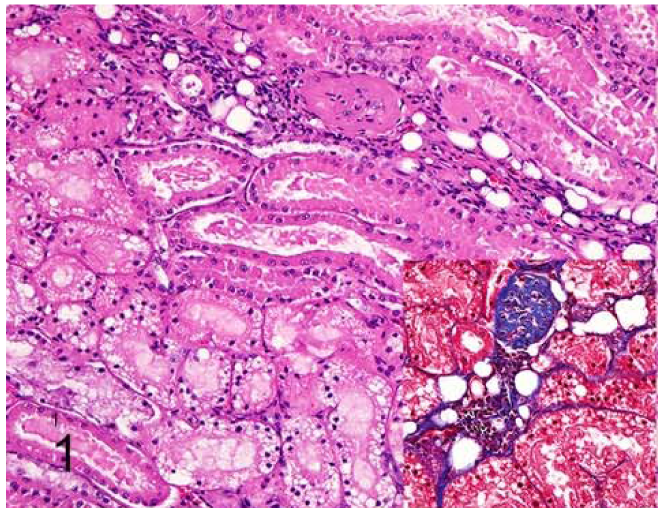
Results from Group 1 (Chronic Kidney Disease, CKD). Groups 2 and 3 not shown. Microscopic appearance (HE, Trichrome). Immunohistochemistry (IHC) using primary antibodies derived from serum of mice co-housed with MKPV infected mice and serum from non-co-housed mice. In Situ Hybridization (ISH) using a probe for Mouse Kidney Parvovirus (MKPV) and Peptidylpropyl Isomerase B (PPIB) and 4-hydroxy-tetrahydrodipicolinate reductase (DapB), which serve as positive and negative control, respectively. Group 1, Case 5. Small amounts (<25%) interstitial inflammation and fibrosis within the cortex with focal or scattered tubular degeneration. HE, Trichrome (inset).

### Immunohistochemistry and RNA *in situ* hybridization

For chromogenic immunohistochemistry and RNA *in situ* hybridization (RNA-ISH), 14-slot (3×5) tissue microarrays (TMAs) were generated with one slot left blank (for orientation purposes). Two 4 mm-diameter tissue punches were obtained at random from the renal cortices in each FFPE block and inserted into the TMA slots. Punches of renal cortex of a normal pig kidney were included as negative controls. In total, four TMA blocks were generated per group, for a total of 12 TMAs. These blocks were sectioned at 5 μm-thickness and processed for RNA-ISH and IHC. For both methods, positive (renal tissue from an NSG mouse with histologic evidence of IBN and positive for MKPV by PCR) and negative (histologically normal renal tissue from an NSG mouse confirmed negative for MKPV by PCR) controls were included.

Chromogenic IHC was performed using previously described mouse sera and methods.^14^ Briefly, slides were incubated with pooled sera derived from three Tac:SW mice co-housed for 3 to 13 weeks with NSG mice shedding MKPV in urine, or with serum from a naïve, non-co-housed Tac:SW, and diluted at 1:1000 dilution. Slides were incubated with the secondary antibody biotin-conjugated horse anti-mouse IgG (Vector Labs, Burlingame CA, USA, Cat #) at 1:500 dilution. Avidin-biotin complete elite was added (part of the Vectastain ABC Elite Kit, Vector Labs, Burlingame, CA, USA), followed by the substrate 3,3’-diaminobenzidine (Sigma, St Louis, MO, USA).^14^

Chromogenic RNA-ISH for MKPV was performed as previously described.^14^ A target probe was designed for a 978-base sequence of MKPV mRNA covering the VP1 and NS1 regions, specifically nucleotides 2444-3421, using RNAscope 2.5 LS Probe (Advanced Cell Diagnostics [ACD], Newark, CA, USA). Feline specific PPIB (ACD, Newark, CA, USA) was used as a positive control and dapB (ACD, Newark, CA, USA) was used as a negative control. Control probes were substituted for the target probe, and adequate hybridization of the controls was confirmed. The completed slides were examined by two ACVP board-certified pathologists (AM, TD).

## Results

Results for groups 1, 2 and 3 are summarized in Table 2. Supplemental Table 1 contains all the relevant signalment and clinicopathologic data. Results for group 1 (CKD cats) are summarized in Figures 1–3. Cats in group 1 had ages ranging from 5-19 years, with a median age of 16 years. Cats in group 2 had ages ranging from 0.2 years to 18 years, with a median age of 6 years. Cats in group 3 had ages ranging from 0.25 to 13 years, with a median age of 6.5 years. All IHC and RNA-ISH positive and negative mouse control tissues showed acceptable positive and negative immunoreactivity and hybridization, respectively. Similarly, MKPV-infected positive control urine produced specific product in PCR reactions using broad-specificity primers 961–962/963 or fechavirus-optimized primers 974-975, with sensitivity for as few as 5 templates (Fig S2–S3). PCR, RNA-ISH and IHC did not reveal the presence of target nucleic acids or antigen in any of the samples we tested (Figs. 4–9). In sections stained with sera of cohoused and non-co-housed mice, brown chromogen was present within approximately 2 μm-diameter round structures within the tubular epithelial cytoplasm interpreted as nonspecific staining (Figs 6–7). Similarly, red chromogen highlighted similar structures when RNA-ISH was performed in addition to diffuse, pinpoint, red chromogen likely representing non-specific staining in the cytoplasm of most samples with no difference between positive (feline PPIB), negative (dapB) and MKPV probes (Figs. 8–9). This staining was distinct from the intense punctate staining expected for a positive reaction with this assay, as observed in sections stained with feline PPIB (figure 9). The intensity of chromogen signal interpreted as non-varied between samples. All samples showed hybridization with feline PPIB probes, confirming adequate preservation of RNA.

**Table 2.** Summary of IHC, ISH and PCR for MKPV performed on three groups of cats. Group 1: cats with CKD; Group 2: immunocompromised cats; Group 3: control cats with no co-morbidities.

| **Group** | **IHC** | **ISH** | **PCR** |
| --- | --- | --- | --- |
| G1 CKD (24) | 24/24 Neg | 24/24 Neg | 24/24 negative for chaphamaparvovirus DNA |
| G2 Immune Suppressed (26) | 26/26 Neg | 26/26 Neg | 1/26 negative for chaphamaparvovirus DNA |
| G3 Control cats (25) | 25/25 Neg | 25/25 Neg | Not Performed |

**Figure 2.**
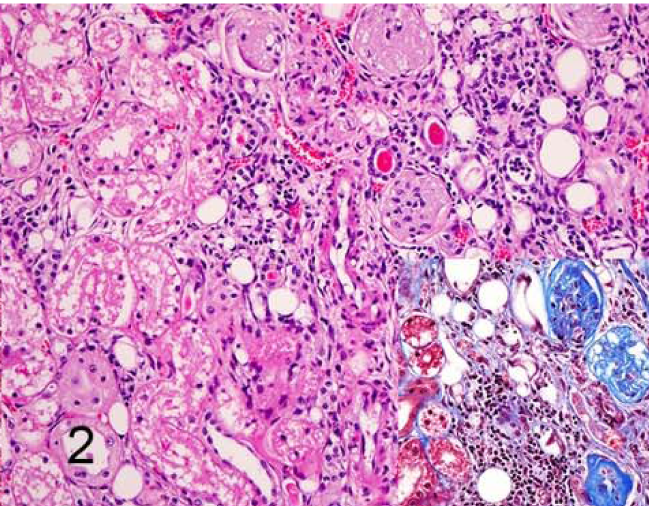
Results from Group 1 (Chronic Kidney Disease, CKD). Groups 2 and 3 not shown. Microscopic appearance (HE, Trichrome). Immunohistochemistry (IHC) using primary antibodies derived from serum of mice co-housed with MKPV infected mice and serum from non-co-housed mice. In Situ Hybridization (ISH) using a probe for Mouse Kidney Parvovirus (MKPV) and Peptidylpropyl Isomerase B (PPIB) and 4-hydroxy-tetrahydrodipicolinate reductase (DapB), which serve as positive and negative control, respectively. Group 1, Case 2. Moderate amounts (25-50%) interstitial inflammation and fibrosis within the cortex with multifocal to coalescing tubular degeneration HE, Trichrome (inset).

**Figure 3.**
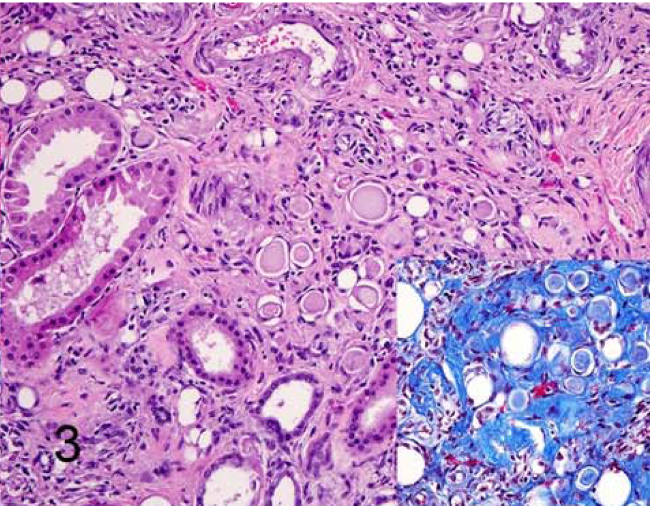
Results from Group 1 (Chronic Kidney Disease, CKD). Groups 2 and 3 not shown. Microscopic appearance (HE, Trichrome). Immunohistochemistry (IHC) using primary antibodies derived from serum of mice co-housed with MKPV infected mice and serum from non-co-housed mice. In Situ Hybridization (ISH) using a probe for Mouse Kidney Parvovirus (MKPV) and Peptidylpropyl Isomerase B (PPIB) and 4-hydroxy-tetrahydrodipicolinate reductase (DapB), which serve as positive and negative control, respectively. Group 1, Case 16. Severe (>50%) interstitial inflammation and fibrosis with tubular degeneration involving entire nephron units. HE, Trichrome (inset).

**Figure 4.**
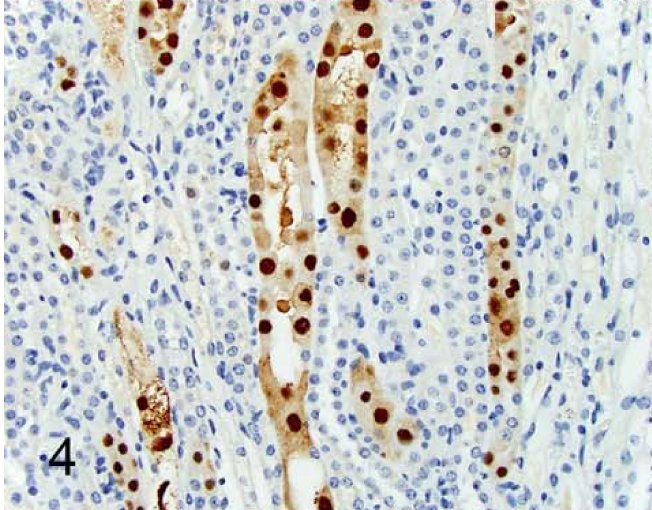
Results from Group 1 (Chronic Kidney Disease, CKD). Groups 2 and 3 not shown. Microscopic appearance (HE, Trichrome). Immunohistochemistry (IHC) using primary antibodies derived from serum of mice co-housed with MKPV infected mice and serum from non-co-housed mice. In Situ Hybridization (ISH) using a probe for Mouse Kidney Parvovirus (MKPV) and Peptidylpropyl Isomerase B (PPIB) and 4-hydroxy-tetrahydrodipicolinate reductase (DapB), which serve as positive and negative control, respectively. Immunohistochemistry with co-housed mouse serum on an NSG mouse infected with MKPV shows strong nuclear and mild cytoplasmic immunoreactivity in tubular cells (positive control).

**Figure 5.**
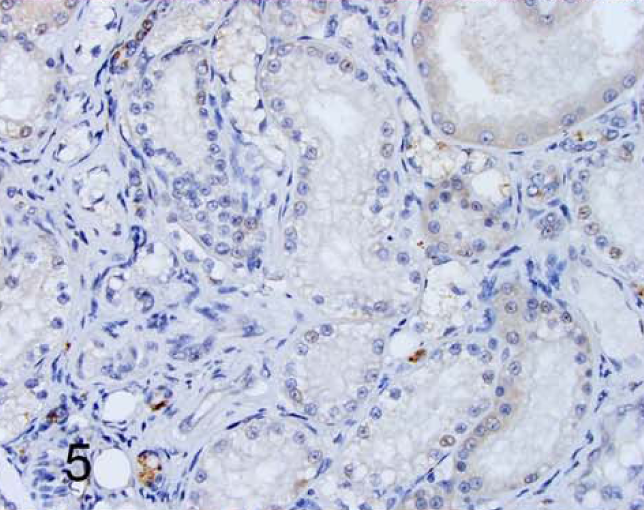
Results from Group 1 (Chronic Kidney Disease, CKD). Groups 2 and 3 not shown. Microscopic appearance (HE, Trichrome). Immunohistochemistry (IHC) using primary antibodies derived from serum of mice co-housed with MKPV infected mice and serum from non-co-housed mice. In Situ Hybridization (ISH) using a probe for Mouse Kidney Parvovirus (MKPV) and Peptidylpropyl Isomerase B (PPIB) and 4-hydroxy-tetrahydrodipicolinate reductase (DapB), which serve as positive and negative control, respectively. Immunohistochemistry with co-housed mouse serum on a kidney sample from group 1 shows weak non-specific background chromogen signal.

**Figure 6.**
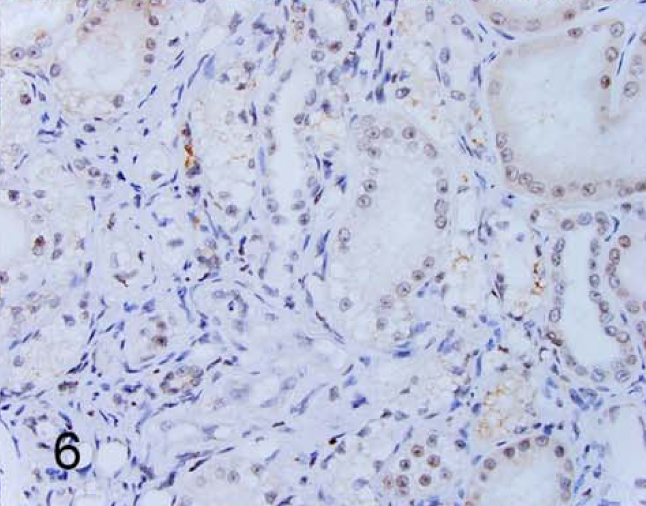
Results from Group 1 (Chronic Kidney Disease, CKD). Groups 2 and 3 not shown. Microscopic appearance (HE, Trichrome). Immunohistochemistry (IHC) using primary antibodies derived from serum of mice co-housed with MKPV infected mice and serum from non-co-housed mice. In Situ Hybridization (ISH) using a probe for Mouse Kidney Parvovirus (MKPV) and Peptidylpropyl Isomerase B (PPIB) and 4-hydroxy-tetrahydrodipicolinate reductase (DapB), which serve as positive and negative control, respectively. Immunohistochemistry with non-co-housed mouse serum on a kidney sample from group 1 shows similar non-specific signal as the co-housed mouse serum assay (negative control).

**Figure 7.**
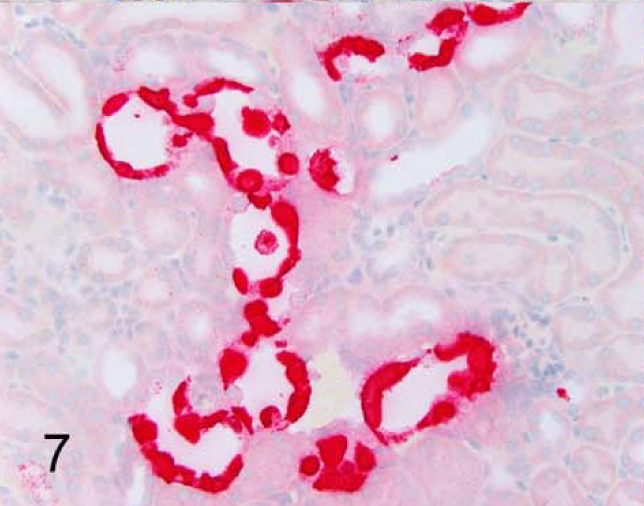
Results from Group 1 (Chronic Kidney Disease, CKD). Groups 2 and 3 not shown. Microscopic appearance (HE, Trichrome). Immunohistochemistry (IHC) using primary antibodies derived from serum of mice co-housed with MKPV infected mice and serum from non-co-housed mice. In Situ Hybridization (ISH) using a probe for Mouse Kidney Parvovirus (MKPV) and Peptidylpropyl Isomerase B (PPIB) and 4-hydroxy-tetrahydrodipicolinate reductase (DapB), which serve as positive and negative control, respectively. In situ hybridization with MKPV probe on an NSG mouse infected with MKPV shows strong nuclear and cytoplasmic hybridization in tubular cells (positive control).

**Figure 8.**
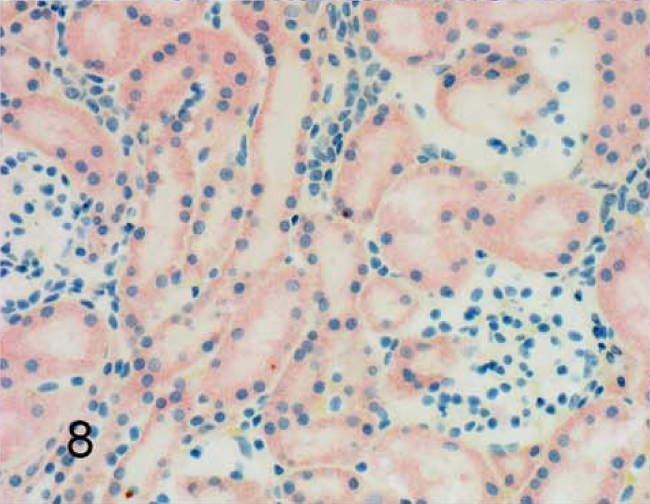
Results from Group 1 (Chronic Kidney Disease, CKD). Groups 2 and 3 not shown. Microscopic appearance (HE, Trichrome). Immunohistochemistry (IHC) using primary antibodies derived from serum of mice co-housed with MKPV infected mice and serum from non-co-housed mice. In Situ Hybridization (ISH) using a probe for Mouse Kidney Parvovirus (MKPV) and Peptidylpropyl Isomerase B (PPIB) and 4-hydroxy-tetrahydrodipicolinate reductase (DapB), which serve as positive and negative control, respectively. In situ hybridization with MKPV probe on a kidney sample from group 1 shows weak non-specific punctate background cytoplasmic chromogen signal.

**Figure 9.**
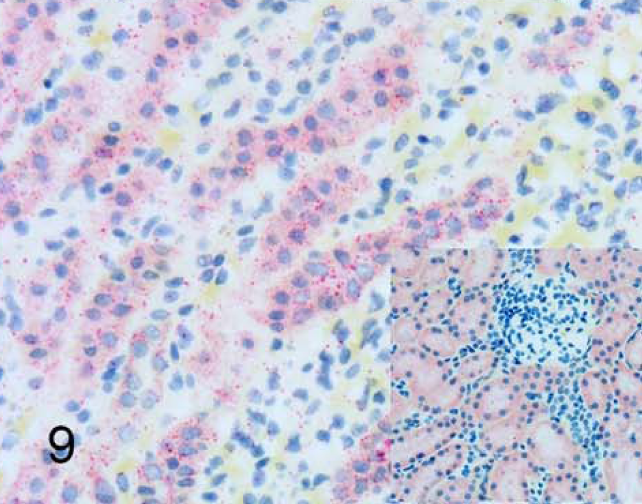
Results from Group 1 (Chronic Kidney Disease, CKD). Groups 2 and 3 not shown. Microscopic appearance (HE, Trichrome). Immunohistochemistry (IHC) using primary antibodies derived from serum of mice co-housed with MKPV infected mice and serum from non-co-housed mice. In Situ Hybridization (ISH) using a probe for Mouse Kidney Parvovirus (MKPV) and Peptidylpropyl Isomerase B (PPIB) and 4-hydroxy-tetrahydrodipicolinate reductase (DapB), which serve as positive and negative control, respectively. In situ hybridization with PPIB shows moderate numbers of cytoplasmic intense punctate dots, consistent with hybridization, indicating good RNA sample quality (positive control). Inset: In situ hybridization with DapB shows weak non-specific punctate background cytoplasmic chromogen signal, similar to MKPV probe (negative control). Magnification 15x objective (insets 40x objective).

## Discussion

MKPV DNA and protein were not detectable in FFPE renal tissues of cats with chronic kidney disease (CKD), cats without renal disease, or immune suppressed cats. As such, both of our hypotheses are incorrect. In addition, from our PCR results, we can likely rule out the recently-discovered cat fechavirus, or a currently unknown but closely related chaphamaparvovirus, as a major cause of chronic kidney disease in domestic cats.^7^

MKPV and MuCPV belong to the novel species *Rodent chaphamaparvovirus 1*, confirmed as the cause of inclusion body nephropathy (IBN) in immune suppressed mice.^14,17^ In laboratory mice, MKPV shows a strong tropism for renal tubular epithelial cells, with documented detectable antigen and RNA in enlarged nuclei (inclusion 14 bodies), achieved with IHC and RNA-ISH, respectively.^14^ Given the widespread distribution of this virus, and concurrent documentation in wild mice from residential buildings in the New York city area, MKPV or MuCPV exposure of domestic cats is very likely.^14,19^ CKD is a common disease of geriatric domestic cats, for which a cause remains elusive. Renal lesions of feline CKD are similar to those of mice infected with MKPV and include cortical tubulointerstitial nephritis, interstitial fibrosis, glomerulosclerosis, hyperplastic arteriolosclerosis and tubular mineralization.^2,14^

The pathogenesis of viral nephropathies are often multifactorial and comprise genetic factors, the renal immunological microenvironment and cellular responses.^13^ Previously-implicated viral causes of feline CKD include Feline Morbillivirus (FmoPV) and Feline Immunodeficiency virus (FIV).^13,16,18,20^ One study found that cats younger than 11 years of age were significantly more likely to be positive for FIV antibodies as 18 compared with cats without CKD.^18^ Reported renal alterations in FIV infected cats encompass predominantly glomerular changes including mesangial cell proliferation and glomerulosclerosis, with a smaller percentage showing immune-mediated glomerulonephritis and amyloidosis.^13^ FmoPV was described with tubulointerstitial nephritis in domestic cats in Hong Kong with detectable antigen observed with IHC in small numbers of cases.^16,20^

Our study is limited by its retrospective nature and usage of FFPE tissues ranging from 2008 to 2018. Antigen, DNA and RNA breakdown over time are possible and formalin fixation may decrease sensitivity, although in the case of RNA-ISH, appropriate positive controls (Feline-PPIB) showed that even the earliest archival tissue showed adequate hybridization. Similarly, the use of TMAs to test all animals may have decreased sensitivity to MKPV but allowed a drastic increase of sample size by decreasing costs 14-fold. In mice, the distribution of MKPV antigen in the kidneys can be multifocal, such that sampling two 4 mm-diameter core biopsies per kidney may be insufficient to detect antigen or RNA.^14^ To account for this, however, thick FFPE scrolls from group 1 were sampled and tested with probes that targeted a well-conserved region of chaphamaparvoviruses, not limited to MKPV. In this study, we used three different approaches to detect viral nucleic acids: RNA-ISH, 40-cycle DNA PCR and chromogenic IHC. Although probably less sensitive than qPCR, which has a detection limit of <10 MKPV genomes, RNA-ISH has higher sensitivity than low cycle number screening PCR, because some mouse FFPE kidney samples negative for MKPV by 25-cycle screening PCR were positive by viral RNA-ISH.^6,14,4^ A disadvantage of RNA-ISH is that variations in viral RNA would likely result in negative hybridization due to the probe’s high specificity. As such, RNA-ISH would only detect MKPV and would be unlikely to detect a related virus. This issue was addressed by PCR from FFPE specimens, using redundant primers designed to recognize several chaphamaparvoviruses previously detected in kidneys, as well as fechavirus, thus covering a broad range of chaphamaparvoviruses. Since DNA is often fragmented or damaged in FFPE samples, PCR sensitivity was maximized by using 40 PCR cycles with a low Tm for the final 30 cycles and by targeting a small amplicon (78bp) with a highly processive polymerase.^6,7^ By definition, the serum from wild-type Tac:SW mice co-housed with MKPV-infected NSG mice is polyclonal. As such, it was expected to recognize multiple epitopes of MKPV, some of which may be conserved among other related chaphamaparvoviruses. However, this presumptive cross-reactivity could not be tested due to the lack of availability of tissues infected by other chaphamaparvoviruses, most of which have been described only by metagenomics analysis without concurrent histopathologic analyses. In addition, the serum would likely recognize any infectious agent to which the Swiss Webster mice have been exposed and that persistently infects any of the cats in our study. While we have clearly demonstrated that this serum can detect MKPV in immunodeficient mice with abundant virus (Fig. 4), the relative sensitivity and specificity of co-housed serum in relation to PCR or ISH is untested.

In our study, we included cats that were affected by CKD (group 1) as well as immunocompromised cats (group 2). In addition, the age range of these 50 veterinary patients was broad (0.2 to 19 years of age). The inclusion of immunocompromised cats reflects a high degree of viral persistence seen in NSG mice, which is likely due to their lack of adaptive immunity. Per our hypothesis, immunocompromised cats may have been at a higher risk of an MKPV infection, although this is evidently not the case. Similarly, including a large age-range in both groups, and of disease severity in the CKD group, decreases the likelihood of false-negative results resulting from a cleared MKPV infection. However, in NSG mice, MKPV antigen and DNA can still be detected in aged mice with severe CKD lesions, and this has more recently been observed in aged immunocompetent Swiss Webster mice with milder lesions as well (A. Michel, unpublished data). Finally, these tissue samples were obtained from a defined geographical location (Northeastern USA) and may not reflect global trends in cats with CKD.

We conclude that MKPV, fechavirus or another chaphamaparvovirus are not a likely cause or contributing factor to CKD in domestic cats and further research into other causative factors should be pursued.

## Author approval

All authors have seen and approved the manuscript. The manuscript has not been accepted or published elsewhere.

## Acknowledgments

This work was funded by the Caspary Research Institute of The Animal Medical Center. AM and SM are supported in part by NCI Cancer Center Support Grant P30 CA008748. TD is supported by the Caspary Research Institute of the Animal Medical Center. BR, LQ and CJ were supported by Australian National Health and Medical Research Council, the Cancer Institute NSW and the Hillcrest Foundation. The authors would like to thank Amanda Ramkissoon for data collection and performing special stains, John D’Allara for TMA preparation, Iveta Simanska for TMA sectioning, and Maria Jiao for performing IHC and RNA-ISH.

---

**Figure S1:**
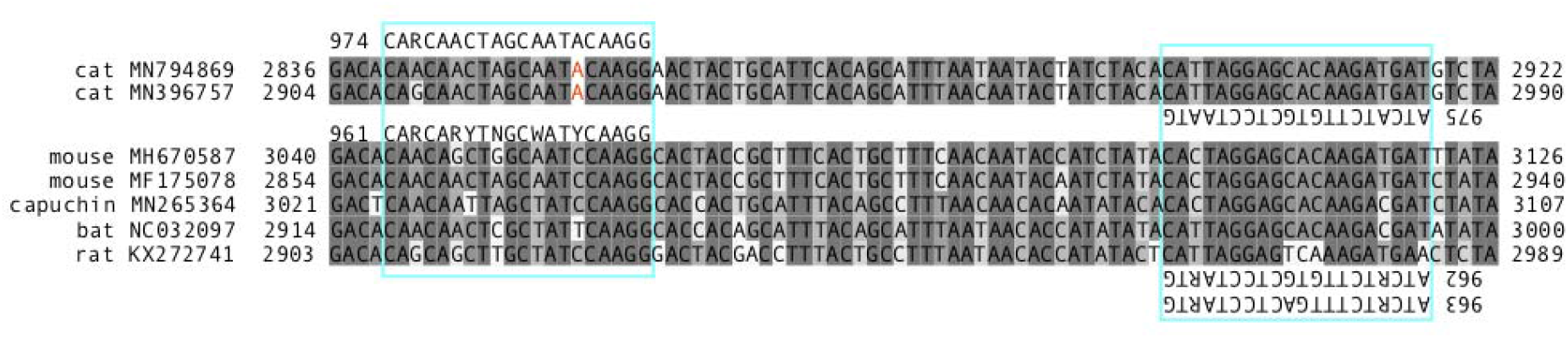
Blue boxes indicate alignment of primers 961, 962, 963, 974 and 975 with VP sequences from several chaphamaparvoviruses - with accession numbers and base positions. A single mismatch between primer 961 and the known cat chaphamaparvoviruses is indicated red.

**Figure S2:**
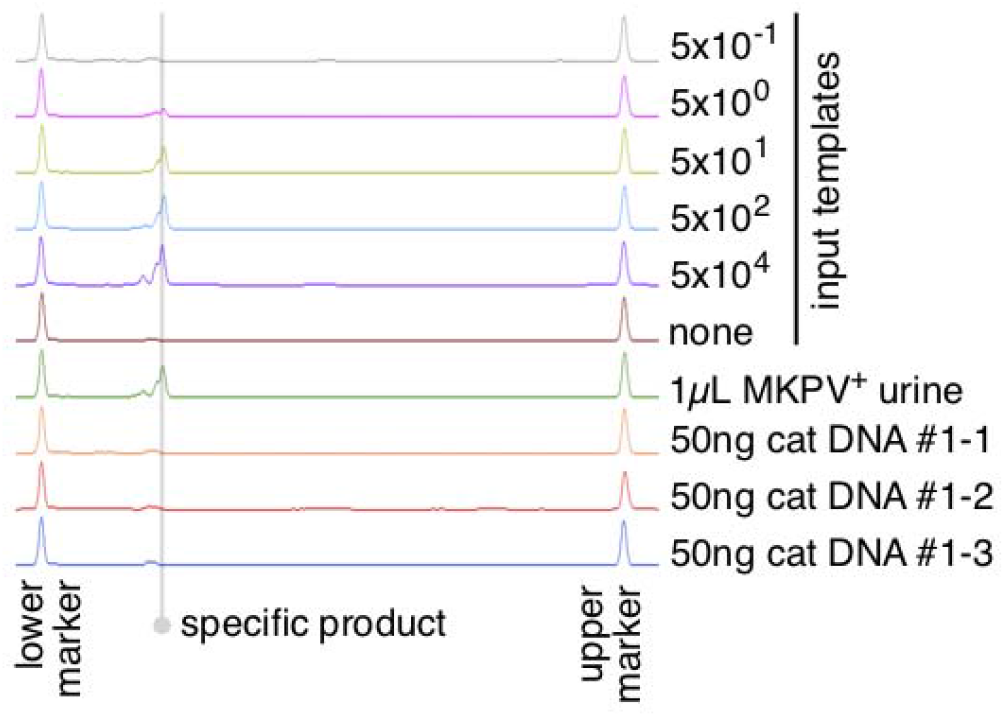
Example capillary electrophoretograms of 40-cycle 974-975 PCR products amplified from a dilution series of oligonucleotide 972 templates, 1μL MKPV-infected mouse urine or 50ng FFPE-extracted cat kidney DNA.

**Figure S3:**
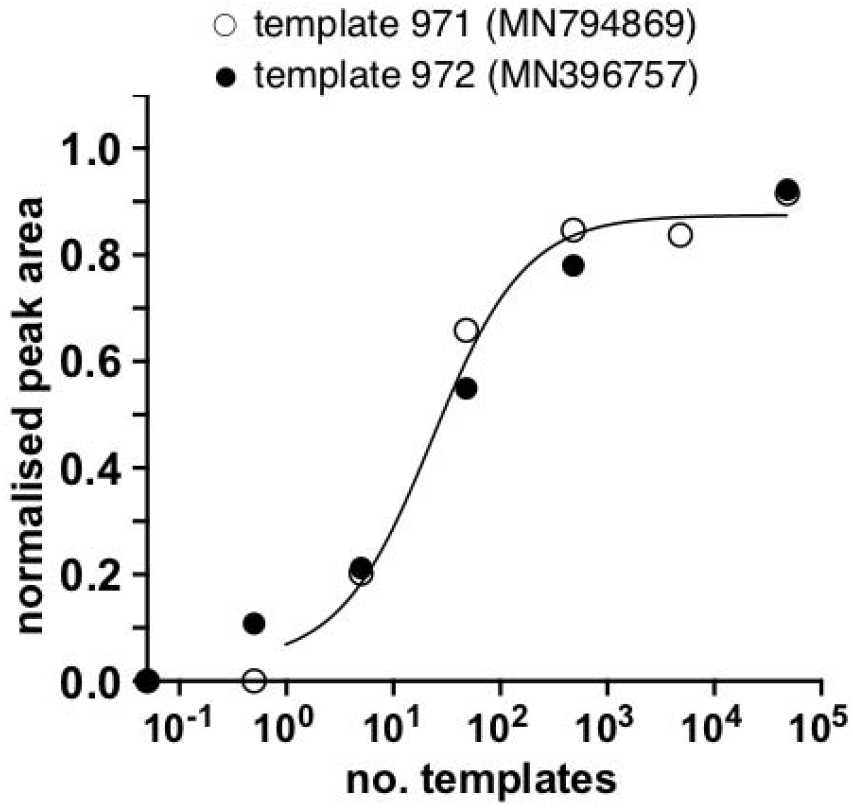
Sensitivity curve (Padé approximant, Prism v8.3.0, GraphPad Software) of normalized peak areas for 40-cycle 974-975 PCR products produced from dilution series of template oligonucleotides 971 or 972.

