## Supplemental Table S1 for "Chaphamaparvovirus Antigen and Nucleic Acids are not Detected in Kidney Tissues from Cats with Chronic Renal Disease or Immunosuppressive Diseases"

| Group | Case | Accession # | Sex | Age | Necropsy date | SDMA (Date Tested) | USG (date tested) | IRIS Stage | IRIS Stage according to Crea only |
|---|---|---|---|---|---|---|---|---|---|
| 1 | 1 | 47816-6 | FS | 16 | 10/6/08 | N/A | 1.000 (6/9/08) | N/A | 3 |
| 1 | 2 | 48003-2 | MC | 17 | 8/13/09 | N/A | 1.015 (8/4/09) | N/A | 2 |
| 1 | 3 | 48060-9 | FS | 8 | 12/7/09 | N/A | 1.018 (12/7/09) | N/A | 2 |
| 1 | 4 | 48105-1 | FS | 7 | 2/22/10 | N/A | 1.020 (11/23/09) | N/A | 3 |
| 1 | 5 | 48248-2 | MC | 16 | 9/7/10 | N/A | 1.012 (5/19/09) | N/A | 2 |
| 1 | 6 | 48363-5 | MC | 17 | 1/11/11 | N/A | 1.014 (12/14/10) | N/A | 4 |
| 1 | 7 | 48398-4 | FS | 16 | 2/24/11 | N/A | 1.018 (10/12/10) | N/A | 2 |
| 1 | 8 | 48566-7 | FS | 19 | 9/19/11 | N/A | 1.020 (9/5/11) | N/A | 2 |
| 1 | 9 | 48624-4 | MC | 17 | 12/26/11 | N/A | 1.020 (12/21/11) | N/A | 4 |
| 1 | 10 | 48807-6 | MC | 17 | 10/26/12 | N/A | N/A | N/A | 3 |
| 1 | 11 | 48863-7 | FS | 5 | 1/15/13 | N/A | 1.019(1/12/13) | N/A | 3 |
| 1 | 12 | 48897-9 | FS | 15 | 2/25/13 | N/A | 1.012 2/18/13 | N/A | 2 |
| 1 | 13 | 48982-1 | MC | 19 | 6/19/13 | N/A | 1.022 (5/2/12) | N/A | 3 |
| 1 | 14 | 49259-7 | MC | 15 | 12/18/15 | N/A | 1.016 (10/29/14) | III | 2 |
| 1 | 15 | 49349-8 | MC | 11 | 5/14/15 | N/A | 1.016 (5/13/15) | N/A | 4 |
| 1 | 16 | 49436-7 | MC | 13 | 9/1/15 | N/A | 1.017 (8/12/15) | II | 3 |
| 1 | 17 | 49486-9 | MC | 15 | 11/23/15 | 29 (11/15/15) | 1.020 (11/15/15) | II | 1 or 2 |
| 1 | 18 | 49612-4 | FS | 16 | 5/20/16 | 26 (5/20/16) | 1.014 (11/16/15) | N/A | 2 |
| 1 | 19 | 49640-1 | FS | 13 | 6/30/16 | N/A | N/A | III | 2 |
| 1 | 20 | 49840-3 | MC | 17 | 3/27/17 | N/A | 1.015 (1/20/17) | II | 2 |
| 1 | 21 | 49937-D | FS | 18 | 8/9/17 | 23 (5/23/17) | 1.017 (5/23/17) | N/A | 2 |
| 1 | 22 | 49967-H | FS | 15 | 10/2/17 | 14 (9/29/17) | 1.017 (9/29/17) | II | 3 |
| 1 | 23 | 50109-D | MC | 19 | 4/2/18 | SDMA 16 (10/3/17) | 1.014 (10/3/17) | N/A | 3 |
| 1 | 24 | 50147-J | MC | 12 | 5/23/18 | SDMA 23 (3/7/18) | N/A | N/A | 2 |
| 1 | 1 | 47814-3 | M | 1 | 10/1/08 | N/A | N/A | N/A | N/A |
| 2 | 2 | 47991-4 | FS | 3 | 7/20/09 | N/A | N/A | N/A | N/A |
| 2 | 3 | 47992-3 | MC | 18 | 7/24/09 | N/A | N/A | N/A | N/A |
| 2 | 4 | 47999-3 | M | 6 | 8/6/09 | N/A | N/A | N/A | N/A |
| 2 | 5 | 48221-4 | FS | 1 | 7/30/10 | N/A | N/A | N/A | N/A |
| 2 | 6 | 48489-8 | MC | 1 | 6/15/11 | N/A | N/A | N/A | N/A |
| 2 | 7 | 48586-3 | MC | 1 | 10/19/11 | N/A | N/A | N/A | N/A |
| 2 | 8 | 48744-1 | FS | 2 | 7/6/12 | N/A | N/A | N/A | N/A |

| | | | | | | | | | |
|---|---|---|---|---|---|---|---|---|---|
| 2 | 9 | 48763-7 | MC | 1 | 7/26/12 | N/A | N/A | N/A | N/A |
| 2 | 10 | 48839-4 | F | 6 | 12/18/12 | N/A | N/A | N/A | N/A |
| 2 | 11 | 48940-2 | FS | 8 | 5/2/13 | N/A | N/A | N/A | N/A |
| 2 | 12 | 48977-8 | MC | 2 | 6/18/13 | N/A | N/A | N/A | N/A |
| 2 | 13 | 49018-11 | MC | 11 | 8/12/13 | N/A | N/A | N/A | N/A |
| 2 | 14 | 49261-6 | MC | 10 | 12/23/14 | N/A | N/A | N/A | N/A |
| 2 | 15 | 49523-2 | F | 6 | 1/7/16 | N/A | N/A | N/A | N/A |
| 2 | 16 | 49667-9 | MC | 1 | 8/11/16 | N/A | N/A | N/A | N/A |
| 2 | 17 | 49696-8 | FS | 8 | 9/19/16 | N/A | N/A | N/A | N/A |
| 2 | 18 | 49733-5 | FS | 12 | 10/26/16 | N/A | N/A | N/A | N/A |
| 2 | 19 | 49747-4 | F | 9 | 11/11/16 | N/A | N/A | N/A | N/A |
| 2 | 20 | 49879-K | MC | 2 | 5/17/17 | N/A | N/A | N/A | N/A |
| 2 | 21 | 49993-H | MC | 12 | 11/6/17 | N/A | N/A | N/A | N/A |
| 2 | 22 | 50078-H | MC | 9 | 2/26/18 | N/A | N/A | N/A | N/A |
| 2 | 23 | 50089-L | FS | 1 | 3/13/18 | N/A | N/A | N/A | N/A |
| 2 | 24 | 50188-J | M | 10 | 7/17/18 | N/A | N/A | N/A | N/A |
| 2 | 25 | 50203-F | FS | 1 | 8/2/18 | N/A | N/A | N/A | N/A |
| 2 | 26 | 48857-6 | MC | 11 | 1/7/13 | N/A | 1.004 (11/20/12) | IV | N/A |
| 3 | 1 | 47804-8 | MC | 8 | 9/15/08 | N/A | N/A | N/A | N/A |
| 3 | 2 | 47910-2 | MC | 3.5 | 3/20/09 | N/A | N/A | N/A | N/A |
| 3 | 3 | 48038-5 | FS | 10 | 10/26/09 | N/A | N/A | N/A | N/A |
| 3 | 4 | 48271-3 | MC | 10 | 10/7/10 | N/A | N/A | N/A | N/A |
| 3 | 5 | 48426-4 | FS | 2 | 3/28/11 | N/A | N/A | N/A | N/A |
| 3 | 6 | 48584-3 | FS | 4 | 10/17/11 | N/A | N/A | N/A | N/A |
| 3 | 7 | 48610-7 | FS | 10 | 11/29/11 | N/A | N/A | N/A | N/A |
| 3 | 8 | 48648-5 | FS | 5 | 1/30/12 | N/A | N/A | N/A | N/A |
| 3 | 9 | 48876-9 | FS | 6 | 1/28/13 | N/A | N/A | N/A | N/A |
| 3 | 10 | 48892-7 | FS | 10 | 2/14/13 | N/A | N/A | N/A | N/A |
| 3 | 11 | 48930-5 | MC | 7 | 4/9/13 | N/A | N/A | N/A | N/A |
| 3 | 12 | 48958-10 | FS | 3 | 5/22/13 | N/A | N/A | N/A | N/A |
| 3 | 13 | 49013-5 | MC | 8 | 8/6/13 | N/A | N/A | N/A | N/A |
| 3 | 14 | 49020-7 | MC | 5 | 8/16/13 | N/A | N/A | N/A | N/A |
| 3 | 15 | 49036-4 | MC | N/A | 9/5/13 | N/A | N/A | N/A | N/A |
| 3 | 16 | 49060-12 | MC | 13 | 10/28/13 | N/A | N/A | N/A | N/A |
| 3 | 17 | 49083-5 | FS | 3 | 12/17/13 | N/A | N/A | N/A | N/A |

| | | | | | | | | | |
|---|---|---|---|---|---|---|---|---|---|
| 3 | 18 | 49114-7 | FS | 10 | 2/19/14 | N/A | N/A | N/A | N/A |
| 3 | 19 | 49179-6 | FS | 7 | 7/7/14 | N/A | N/A | N/A | N/A |
| 3 | 20 | 49191-11 | M | 13 | 8/4/14 | N/A | N/A | N/A | N/A |
| 3 | 21 | 49300-3 | FS | 5 | 2/19/15 | N/A | N/A | N/A | N/A |
| 3 | 22 | 49458-7 | FS | 12 | 10/5/15 | 10 (9/14/15) | N/A | N/A | N/A |
| 3 | 23 | 49479-8 | MC | 4 | 11/6/15 | N/A | N/A | N/A | N/A |
| 3 | 24 | 49883-E | FS | 2 | 5/22/17 | N/A | N/A | N/A | N/A |
| 3 | 25 | 50095-G | MC | 6 | 3/19/18 | N/A | N/A | N/A | N/A |

SDMA: Symmetric dimethylarginine (Reference range: 0-14)
BUN: Blood urea nitrogen (Reference range: 16-37 mg/dL)
Crea: Creatinine (Reference range: 0.9-2.3 mg/dL)
USG: Urine specific gravity (Reference range: <1.035)
FS: Female spayed
MC: Male castrated
F: Female
M: Male

| Diagnosed immunosuppressive disease | BUN/Creat (Date Tested) | Inflammation cortex 0-3 | Fibrosis cortex 0-3 | Tubular Degeneration 0-3 | Total | Normal (1-2), mild (3-4), moderate (5-6) and severe (7-9) |
|---|---|---|---|---|---|---|
| None | 74/4.2 (9/29/08) | 1 | 2 | 1 | 4 | Mild |
| None | 40/2.1 (7/28/09) | 2 | 2 | 2 | 6 | Moderate |
| None | 88/1.9 (12/5/09) | 3 | 2 | 2 | 7 | Severe |
| None | 85/3.1 (1/25/09) | 2 | 3 | 3 | 8 | Severe |
| None | 51/1.9 (8/2/10) | 1 | 1 | 1 | 3 | Mild |
| None | 131/5.5 (12/14/10) | 2 | 2 | 2 | 6 | Moderate |
| None | 39/2.6 (2/23/11) | 1 | 1 | 1 | 3 | Mild |
| None | 53/1.6 (9/5/11) | 1 | 2 | 1 | 4 | Mild |
| None | 106/6.3 (12/22/11) | 3 | 3 | 3 | 9 | Severe |
| None | 73/4.5 (10/25/12) | 2 | 2 | 2 | 6 | Moderate |
| None | 133/4.2 (1/15/13) | 2 | 2 | 2 | 6 | Moderate |
| None | 80/2.0 (2/18/13) | 1 | 2 | 2 | 5 | Moderate |
| None | 50/4.0 (5/2/12) | 3 | 3 | 3 | 9 | Severe |
| None | 41/2.3 (10/29/14) | 2 | 2 | 3 | 7 | Severe |
| None | 194/6.7 (5/13/15) | 2 | 2 | 2 | 6 | Moderate |
| None | 111/3.9 (7/29/15) | 2 | 3 | 3 | 8 | Severe |
| None | 44/1.5 (11/15/15) | 2 | 1 | 1 | 4 | Mild |
| None | 130/2.1 (5/20/16) | 3 | 3 | 3 | 9 | Severe |
| None | 44/2.3 (6/29/16) | 1 | 1 | 1 | 3 | Mild |
| None | 38/1.9 (3/13/17) | 2 | 2 | 2 | 6 | Moderate |
| None | 94/2.3 (5/23/17) | 1 | 2 | 1 | 4 | Mild |
| None | 45/2.9 (9/29/17) | 1 | 2 | 1 | 4 | Mild |
| None | 74/3.8 (10/3/17) | 2 | 2 | 3 | 7 | Severe |
| None | 45/2.2 (3/7/18) | 2 | 1 | 2 | 5 | Moderate |
| FeLv | N/A | 0 | 0 | 0 | 0 | Normal |
| FeLv | N/A | 0 | 0 | 1 | 1 | Mild |
| FIV | N/A | 1 | 1 | 1 | 3 | Mild |
| Panleuk | N/A | 0 | 0 | 0 | 0 | Normal |
| Panleuk | N/A | 0 | 0 | 0 | 0 | Normal |
| FIPV | N/A | 1 | 0 | 0 | 1 | Mild |
| FIPV | N/A | 2 | 0 | 2 | 4 | Mild |
| FeLV , FIPV | N/A | 1 | 0 | 2 | 3 | Mild |

| | | | | | | |
|---|---|---|---|---|---|---|
| FIPV | N/A | 3 | 0 | 2 | 5 | Moderate |
| FIPV | N/A | | | | | |
| FIPV | N/A | 1 | 0 | 0 | 1 | Mild |
| FeLV | 17/1.6 (6/12/13) | 0 | 1 | 0 | 1 | Mild |
| FIV | N/A | 2 | 1 | 3 | 5 | Moderate |
| FIV | N/A | 2 | 2 | 1 | 5 | Moderate |
| FIPV | N/A | 0 | 0 | 0 | 0 | Normal |
| FIPV | N/A | 1 | 0 | 2 | 3 | Mild |
| Cryptococcus | N/A | 0 | 0 | 0 | 0 | Normal |
| FHV-1 | N/A | 1 | 1 | 2 | 4 | Mild |
| Panleuk | N/A | 0 | 0 | 0 | 0 | Normal |
| FeLV | N/A | 0 | 0 | 0 | 0 | Normal |
| FIV | N/A | 1 | 1 | 0 | 2 | Mild |
| FHV-1 | N/A | 1 | 0 | 1 | 2 | Mild |
| FIPV | N/A | 1 | 1 | 1 | 3 | Mild |
| FIPV | N/A | 1 | 0 | 1 | 2 | Mild |
| FeLV | N/A | 1 | 0 | 0 | 1 | Mild |
| Cryptococcus | 49/3.1 (11/20/12) | 2 | 2 | 3 | 7 | Severe |
| None | 29 / 1.2 (8/7/08) | 0 | 0 | 0 | 0 | Normal |
| None | 22/0.4 (3/19/09) | 0 | 0 | 0 | 0 | Normal |
| None | 31/1.9 (10/23/09) | 0 | 0 | 0 | 0 | Normal |
| None | 32/0.7 (10/6/2010) | 0 | 0 | 0 | 0 | Normal |
| None | 25/0.5 (3/27/11) | 0 | 0 | 0 | 0 | Normal |
| None | 17/1.3 (6/9/11) | 0 | 0 | 0 | 0 | Normal |
| None | 19/1.2 (10/27/11) | 0 | 0 | 0 | 0 | Normal |
| None | 22/1.1 (11/18/2010) | 0 | 0 | 0 | 0 | Normal |
| None | 20/1.1 (12/28/12) | 0 | 0 | 0 | 0 | Normal |
| None | 24/0.7 (2/12/13) | 0 | 0 | 0 | 0 | Normal |
| None | 16/1.5 (4/8/13) | 0 | 0 | 0 | 0 | Normal |
| None | 16/1.3 (5/21/13) | 0 | 0 | 0 | 0 | Normal |
| None | 14/0.9 (8/3/13) | 0 | 0 | 0 | 0 | Normal |
| None | 20/1.7 (8/14/13) | 0 | 0 | 0 | 0 | Normal |
| None | 16/0.9 (9/2/13) | 0 | 0 | 0 | 0 | Normal |
| None | 30/1.4 (10/18/13) | 0 | 0 | 0 | 0 | Normal |
| None | 15/1.5 (12/12/13) | 0 | 0 | 0 | 0 | Normal |

| | | | | | | |
|---|---|---|---|---|---|---|
| None | 20/1.2 (2/8/14) | 0 | 0 | 0 | 0 | Normal |
| None | 15/1.1 (7/1/14) | 0 | 0 | 0 | 0 | Normal |
| None | 19/0.4 (7/31/14) | 0 | 0 | 0 | 0 | Normal |
| None | 37/0.8 (2/17/15) | 0 | 0 | 0 | 0 | Normal |
| None | 22/1.4 (9/14/15) | 0 | 0 | 0 | 0 | Normal |
| None | 18/0.7 (11/4/15) | 0 | 0 | 0 | 0 | Normal |
| None | 27/1.0 (4/25/17) | 0 | 0 | 0 | 0 | Normal |
| None | 19/1.5 (3/12/17) | 0 | 0 | 0 | 0 | Normal |
